# The *Gossypium anomalum* genome as a resource for cotton improvement and evolutionary analysis of hybrid incompatibility

**DOI:** 10.1101/2021.06.16.448676

**Authors:** Corrinne E. Grover, Daojun Yuan, Mark A. Arick, Emma R. Miller, Guanjing Hu, Daniel G. Peterson, Jonathan F. Wendel, Joshua A. Udall

## Abstract

Cotton is an important crop that has been the beneficiary of multiple genome sequencing efforts, including diverse representatives of wild species for germplasm development. *Gossypium anomalum* is a wild African diploid species that harbors stress-resistance and fiber-related traits with potential application to modern breeding efforts. In addition, this species is a natural source of cytoplasmic male sterility and a resource for understanding hybrid lethality in the genus. Here we report a high-quality *de novo* genome assembly for *G. anomalum* and characterize this genome relative to existing genome sequences in cotton. In addition, we use the synthetic allopolyploids 2(A2D1) and 2(A2D3) to discover regions in the *G. anomalum* genome potentially involved in hybrid lethality, a possibility enabled by introgression of regions homologous to the D3 (*G. davidsonii*) lethality loci into the synthetic 2(A2D3) allopolyploid.

## Introduction

The genus *Gossypium* is responsible for providing a majority of natural textile fiber through cultivation of its four independently domesticated species. Recent efforts in genome sequencing have resulted in high-quality genomes for all domesticated species (Yuan *et al*. 2015; Chen *et al*. 2020; Huang *et al*. 2020) and for other important species (Paterson *et al*. 2012; Udall *et al*. 2019; Chen *et al*. 2020). Recent efforts at sequencing additional wild cotton species (Udall *et al*. 2019; Grover *et al*. 2020, 2021) have resulted in several high-quality resources for exploring the evolution of agronomically favorable traits, e.g., stress resistance, that are found naturally in the wild cotton species.

Comprising more than 50 known species, the diploid species of cotton have been placed into genome groups (known as A-G, and K) based on meiotic chromosome associations and sequence similarities (see Wang et al. 2018 for review). The wild African species *G. anomalum* Waw. & Peyr. is one of four species comprising the “B-genome” cottons. The B-genome cottons are *G. anomalum* (B1), *G. triphyllum* (B2), *G. capitis-viridis* (B3), and perhaps the poorly understood *G. trifurcatum* (Vollesen 1987; Fryxell 1992; Wendel *et al*. 2010), although relationships for the latter, rare species are unclear (Wang et al., 2018). All of these species are in clades that are close relatives of the diploid domesticated species *G. arboreum* and *G. herbaceum. Gossypium anomalum* has a large but disjunct geographic range, encompassing southwest Africa, centered in Namibia (*G. anomalum* subsp. *anomalum*), and then also a broad distribution in northern Africa (*G. anomalum* subsp. *Senarense*; Vollesen 1987; Fryxell 1992). Although the species has no obvious traits of agronomic interest, *G. anomalum* has many understudied characteristics that may be useful in breeding programs and understanding the evolution of favorable phenotypes.

The fiber of *G. anomalum* is short and not spinnable, but *G. anomalum* has been considered a potential source for fiber fineness and strength (Mehetre 2010), and the xerophytic nature of *G. anomalum* makes it a candidate for understanding drought resistance in cotton species. *Gossypium anomalum* also exhibits natural resistance to various cotton pests, including jassids (Mammadov *et al*. 2018), bacterial blight/blackarm (Knight 1954; Fryxell *et al*. 1984), mites (Mehetre 2010), bollworms (Mehetre 2010), and rust (Fryxell *et al*. 1984; Mehetre 2010). Mechanisms underlying pest resistance are understudied, but it is clear that investigation of the *G. anomalum* genome may illuminate valuable genes and alleles underlying resistance (Fryxell *et al*. 1984), as demonstrated by hybridization experiments involving *G. anomalum* and cultivated cottons (Mehetre 2010).

In addition to stress resistance and fiber quality traits, *G. anomalum* is both a source of cytoplasmic male sterility (Meyer and Meyer 1965; Marshall *et al*. 1974) and one of the few cotton species that can be crossed with cottons from subsection *Integrifolia* (i.e., *G. klotzschianum* and *G. davidsonii*; (Hutchinson *et al*. 1947)), which generally exhibit hybrid lethality in other crosses. Both cytoplasmic male sterility and *Integrifolia* derived lethality have applications in cotton (Weaver and Weaver 1977; Lee 1981a; Stelly *et al*. 1988; Stelly 1990; Suzuki *et al*. 2013; Bohra *et al*. 2016), the latter being accessible only in crosses that are non-lethal, e.g., with *G. anomalum*.

Here we describe a high-quality, *de novo* genome assembly for *G. anomalum*, the first for a member of *Gossypium* section *Anomala*, which are colloquially known as the “B-genome” cottons (Wang *et al*. 2018). This genome provides a genetic repository for investigating potentially valuable agronomic traits.

## Methods & Materials

### Plant material and sequencing methods

*Gossypium anomalum* was grown from seed under greenhouse conditions at Brigham Young University (BYU), and mature leaves were collected for sequencing. High-quality DNA was extracted via CTAB (Kidwell and Osborn 1992) and subsequently quantified using a Qubit Fluorometer (ThermoFisher, Inc.). DNA was size selected for fragments >18 kb on the BluePippen (Sage Science, LLC) prior to library construction; fragment size was verified using a Fragment Analyzer (Advanced Analytical Technologies, Inc). A single PacBio (Pacific Biosciences) sequencing library was constructed by the BYU DNA Sequencing Center (DNASC), and 15 PacBio cells were sequenced using the Sequel system. Raw reads were assembled using Canu V1.4 with default parameters (Koren *et al*. 2017).

Leaf tissue was shipped to PhaseGenomics LLC for DNA extraction and HiC library construction. HiC libraries were sequenced on the Illumina HiSeq 2500 (PE125 bp) at the BYU DNASC, and the resulting reads were used to join contigs. JuiceBox (Durand et al., 2016) was used in conjunction with the HiC reads to correct the assembly based on the association frequency between paired-ends. A custom python script (available through PhaseGenomics, LLC) was used to construct the final genome sequence of *G. anomalum*, which consists of 13 scaffolds corresponding to the haploid complement of chromosomes.

### Repeat and gene annotation

Repeats were identified using RepeatMasker (Smit *et al*. 2015) and a custom library consisting of Repbase 23.04 repeats (Bao *et al*. 2015) with cotton-specific repeats (Grover *et al*. 2020). RepeatMasker run parameters were set to a high-sensitivity scan that only masked transposable elements (TEs). Multiple hits were aggregated using “One code to find them all” using default parameters (Bailly-Bechet *et al*. 2014), and the resulting output was summarized in R/4.0.3 (R Core Team 2020) using *dplyr* /2.0.0 (Wickham *et al*. 2015). Repeats were quantified relative to other cotton species with resequencing data downloaded from the GenBank Short-Read Archive (Supplementary Table 1) using the RepeatExplorer pipeline (Novák *et al*. 2010), and results were parsed in R/4.0.3 (R Core Team 2020) as previously described (Grover *et al*. 2020). All code is available at https://github.com/Wendellab/anomalum.

Genome annotations were conducted using existing RNA-seq data from tissues of closely related species (Supplementary Table 2) as previously described (Grover *et al*. 2021). Hisat2 was used to map each RNA-seq library to the hard-masked *G. anomalum* genome [v2.1.0] (Kim *et al*. 2015), and *de novo* transcriptome assemblies were generated via StringTie [v2.1.1] (Pertea *et al*. 2015) and Cufflinks [v2.2.1] (Ghosh and Chan 2016). These RNA-seq assemblies were combined with a Trinity [v2.8.6] (Grabherr *et al*. 2011) reference-guided transcriptome assembly and splice junction information from Portcullis [v1.2.2] (Mapleson *et al*. 2018) in Mikado [v1.2.4] (Venturini *et al*. 2018). GeneMark [v4.38] (Borodovsky and Lomsadze 2011) generated annotations were used in BRAKER2 [v2.1.2] (Hoff *et al*. 2019) to train Augustus [v3.3.2] (Stanke *et al*. 2006). MAKER2 [v2.31.10] (Holt and Yandell 2011; Campbell *et al*. 2014) integrated gene predictions from all three sources, i.e., BRAKER2 trained Augustus, GeneMark, and Mikado, with additional evidence from all available *Gossypium* ESTs (NCBI nt database with the filters “txid3633” and “is_est”), all curated proteins in Uniprot Swissprot [v2019_07] (UniProt Consortium 2008), and all annotated proteins from the *G. hirsutum* (https://www.cottongen.org/species/Gossypium_hirsutum/jgi-AD1_genome_v1.1) and *G. raimondii* (Paterson *et al*. 2012) genomes.

Each gene model was scored by Maker using the annotation edit distance (AED -(Eilbeck *et al*. 2009; Holt and Yandell 2011; Yandell and Ence 2012) relative to EST and protein evidence, and gene models with an AED less than 0.37 were retained. These gene models were functionally annotated using InterProScan [v5.47-82.0] (Jones *et al*. 2014) and BlastP [v2.9.0+] (Camacho *et al*. 2009) searches against the Uniprot SwissProt database. Orthologous relationships between *G. anomalum* and other sequenced diploid cotton genomes, i.e., *G. longicalyx* (Grover et al. 2020), *G. arboreum* (Du *et al*. 2018), *G. herbaceum (Huang et al. 2020), G. raimondii* (Paterson *et al*. 2012) are derived from previously published (Grover *et al*. 2021) OrthoFinder analyses (Emms and Kelly 2015, 2019). All genomes are hosted through CottonGen (https://www.cottongen.org; (Yu *et al*. 2014) and running parameters are available from https://github.com/Wendellab/anomalum.

### *G. anomalum* introgression in the synthetic allotetraploid, 2(A2D3)

A synthetic allotetraploid, i.e., 2(A2D3), was generated by Joshua Lee in the late 1970s to early 1980s. The first step in producing the allotetraploid involved crossing *G. anomalum* (B1) with *G. arboreum* (A2; Supplementary Figure 2). The latter species is incompatible with *G. davidsonii* (D3), as are all species tested except *G. anomalum; G. anomalum* likely possesses a null allele for the lethality locus. By repeatedly backcrossing the *G. anomalum* compatibility region into the recipient *G. arboreum* parent, Lee was able to create a *G. arboreum* line that was compatible with *G. davidsonii*. This was subsequently used to generate a diploid hybrid with *G. davidsonii*, i.e., A2 x D3. Subsequent doubling of this hybrid generated the synthetic 2(A2D3), and this plant has been subsequently maintained by Jonathan Wendel in the Iowa State University greenhouse since the mid-1980s. We downloaded previously generated reads from this synthetic allotetraploid (Supplementary Table 1), along with reads for an additional synthetic allotetraploid, 2(A2D1) (Beasley 1940). Chromosomes from all three diploid species, i.e., *G. arboreum, G. anomalum*, and *G. davidsonii*, were combined to generate an *in silico* genome designated “ABD”. Mapping of the 2(A2D3) reads to the ABD genome identified reads which best match *G. anomalum*. To verify the mapping results, we also mapped reads from *G. arboreum, G. davidsonii*, and the 2(A2D1) synthetic to the same (synthetic) ABD genome. The 2(A2D1) synthetic was included as an additional control because the *G. arboreum* (A2) used in this initial cross (i.e., *var. neglectum* (Beasley 1940)) did not include known introgression from other cotton species. All reads were mapped to the ABD genome using BWA [v0.7.17] (Li and Durbin 2009), and samtools [v1.9] (Li *et al*. 2009) was used to select the reads from each species that uniquely mapped (mapq >= 30) to the *G. anomalum* genome. Contiguous regions of uniquely mapped reads were combined in bedtools [v2.28.0] (Quinlan 2014) for each of the control libraries, i.e., *G. arboreum, G. davidsonii*, and 2(A2D1) to identify putative regions of ambiguity (*i*.*e*., where reads may preferentially map to the *G. anomalum* chromosomes by chance). Overlapping regions between the mapping results of 2(A2D1) and 2(A2D3) were filtered to retain regions where only 2(A2D3) reads mapped to the *G. anomalum* genome sequence. Regions < 5 kb in length were then discarded. These filters resulted in a high-confidence set of reads that were likely derived from the *G. anomalum* introgression specific to 2(A2D3).

## Data availability

The *G. anomalum* genome sequence and raw data are available at NCBI under PRJNA421337 and CottonGen (https://www.cottongen.org/). Supplemental files are available from figshare.

## Results and Discussion

### Genome assembly and annotation

Here we report a *de novo* genome assembly for *G. anomalum* using 55x coverage of PacBio reads and 140.5 million (M) HiC reads. The initial assembly yielded 229 contigs with an N50 of 11 Mb. HiC information was integrated to produce a more contiguous assembly, consisting of 13 chromosomes with an average length of 92 Mb and containing only 20.7 kb (0.002%) gap sequence within the chromosomal scaffolds. The total assembly length is 88% of the estimated 1359 Mb genome (Hendrix and Stewart 2005).

BUSCO analysis (Waterhouse *et al*. 2017) of the 13 assembled chromosomes recovered 97.1% complete BUSCOs from the 2326 BUSCO groups comprising euidcots_odb10 database (Table 1). In general, most BUSCOs were both complete and single copy (89.5%), with a low level of duplication (7.6%). Few BUSCOs were fragmented (0.5%) or missing (2.4%), which indicates a general completeness of the assembly. Dotplot comparisons to other cotton genomes (Figure 1) further confirms that the *G. anomalum* assembly is similar to or superior to recently published genomes.

**Table 1.**
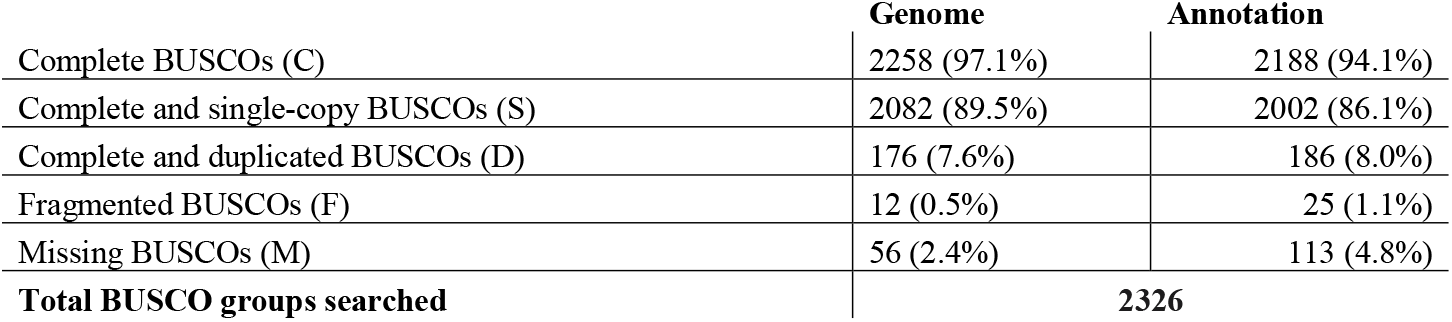
BUSCO scores for the genome and transcriptome of *G. anomalum*.

|  | Genome | Annotation |
| --- | --- | --- |
| Complete BUSCOs (C) | 2258 (97.1%) | 2188 (94.1%) |
| Complete and single-copy BUSCOs (S) | 2082 (89.5%) | 2002 (86.1%) |
| Complete and duplicated BUSCOs (D) | 176 (7.6%) | 186 (8.0%) |
| Fragmented BUSCOs (F) | 12 (0.5%) | 25 (1.1%) |
| Missing BUSCOs (M) | 56 (2.4%) | 113 (4.8%) |
| **Total BUSCO groups searched** | **2326** |  |

**Figure 1.**
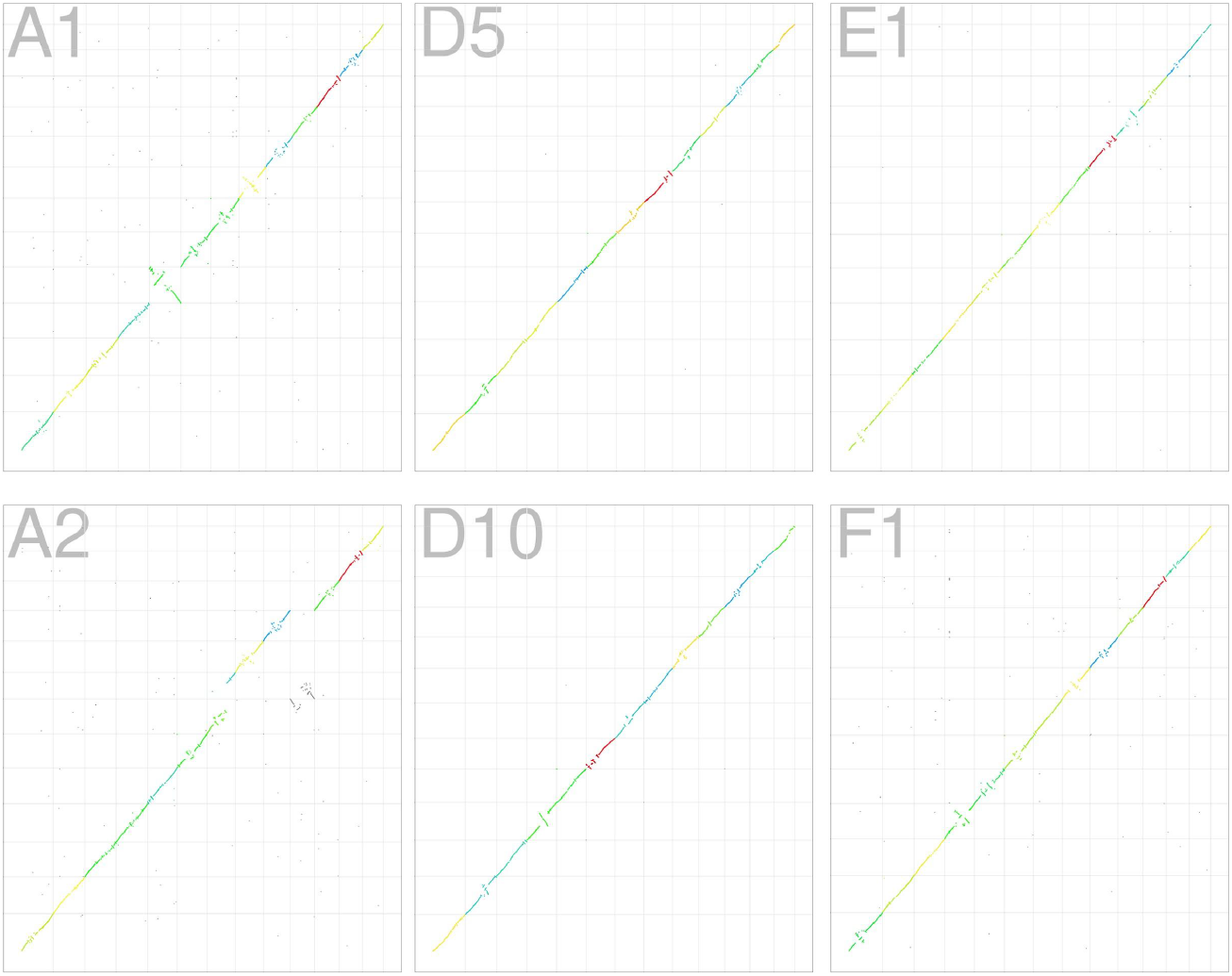
Pairwise comparisons of *G. anomalum* with *G. herbaceum* (A1; (Huang *et al*. 2020), *G. raimondii* (D5; (Udall *et al*. 2019), *G. stocksii (Grover et al. 2021), G. arboreum* (A2; (Huang *et al*. 2020), *G. turneri* (D10; (Udall *et al*. 2019), and *G. longicalyx* (F1; (Grover *et al*. 2020).

Genome annotation produced 37,830 primary transcripts, which is similar to other cotton diploids (Paterson *et al*. 2012; Du *et al*. 2018; Udall *et al*. 2019; Grover *et al*. 2020, 2021; Huang *et al*. 2020; Wang *et al*. 2021) whose gene numbers range between 34,928 (Grover *et al*. 2021) in *G. stocksii* to 43,952 (Huang *et al*. 2020) in *G. herbaceum*. BUSCO analysis of the transcriptome exhibited similar quality to the genome, with 86.1% complete and single copy and few duplicated or missing (8% and 4.8%, respectively). Ortholog analysis of primary transcripts suggests that the pattern of orthogroups including *G. anomalum* is similar to other diploid cotton species, although the number of genes not assigned to orthogroups is fewer than previously noted (Grover *et al*. 2021), whereas the number of species-specific orthogroups is higher, albeit still low (Table 2; Supplementary Table 3).

**Table 2:**
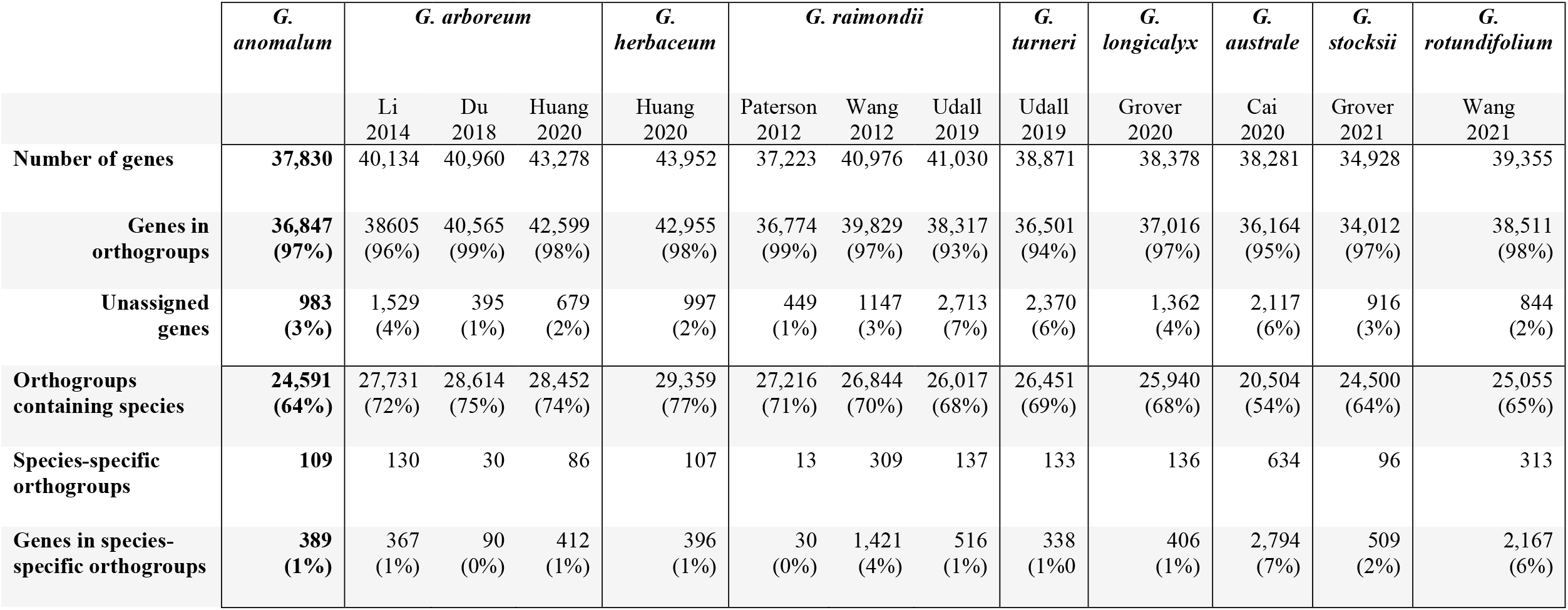
Orthogroup relationships between *G. anomalum* and other cotton diploid genomes (primary transcripts only).

|  | *G.*<br>*anomalum* | *G. arboreum* |  |  | *G.*<br>*herbaceum* | *G. raimondii* |  |  | *G.*<br>*turneri* | *G.*<br>*longicalyx* | *G.*<br>*australe* | *G.*<br>*stocksii* | *G.*<br>*rotundifolium* |
| --- | --- | --- | --- | --- | --- | --- | --- | --- | --- | --- | --- | --- | --- |
|  |  | Li<br>2014 | Du<br>2018 | Huang<br>2020 | Huang<br>2020 | Paterson<br>2012 | Wang<br>2012 | Udall<br>2019 | Udall<br>2019 | Grover<br>2020 | Cai<br>2020 | Grover<br>2021 | Wang<br>2021 |
| **Number of genes** | **37,830** | 40,134 | 40,960 | 43,278 | 43,952 | 37,223 | 40,976 | 41,030 | 38,871 | 38,378 | 38,281 | 34,928 | 39,355 |
| **Genes in orthogroups** | **36,847 (97%)** | 38605 (96%) | 40,565 (99%) | 42,599 (98%) | 42,955 (98%) | 36,774 (99%) | 39,829 (97%) | 38,317 (93%) | 36,501 (94%) | 37,016 (97%) | 36,164 (95%) | 34,012 (97%) | 38,511 (98%) |
| **Unassigned genes** | **983 (3%)** | 1,529 (4%) | 395 (1%) | 679 (2%) | 997 (2%) | 449 (1%) | 1147 (3%) | 2,713 (7%) | 2,370 (6%) | 1,362 (4%) | 2,117 (6%) | 916 (3%) | 844 (2%) |
| **Orthogroups containing species** | **24,591 (64%)** | 27,731 (72%) | 28,614 (75%) | 28,452 (74%) | 29,359 (77%) | 27,216 (71%) | 26,844 (70%) | 26,017 (68%) | 26,451 (69%) | 25,940 (68%) | 20,504 (54%) | 24,500 (64%) | 25,055 (65%) |
| **Species-specific orthogroups** | **109** | 130 | 30 | 86 | 107 | 13 | 309 | 137 | 133 | 136 | 634 | 96 | 313 |
| **Genes in species-specific orthogroups** | **389 (1%)** | 367 (1%) | 90 (0%) | 412 (1%) | 396 (1%) | 30 (0%) | 1,421 (4%) | 516 (1%) | 338 (1%0 | 406 (1%) | 2,794 (7%) | 509 (2%) | 2,167 (6%) |

### Repeats

Both *de novo* TE prediction (Bailly-Bechet *et al*. 2014; Smit *et al*. 2015) and repetitive clustering (Novák *et al*. 2010) were used to assess repetitive elements in the *G. anomalum* genome. As with *G. longicalyx* (Grover *et al*. 2020), RepeatExplorer estimated a larger proportion of the *G. anomalum* genome as repetitive (46.5%) compared to RepeatMasker (42%). Estimates for the different TE categories surveyed (e.g., DNA, Ty3/*gypsy*, Ty1/*copia*, etc.) were generally consistent between the two methods (Supplementary Table 4), although RepeatMasker recovered far more *copia* elements than did RepeatExplorer (47.7 Mbp, versus 29.1 Mbp). This is likely due to the inability of RepeatExplorer to efficiently categorize *copia*-like elements in this genome, instead placing them in a general “LTR” category (21.9 Mbp, versus 0 Mbp for RepeatMasker). As is common among plants, most of the repetitive sequence recovered by both methods was attributed to *gypsy* elements, which occupy 38% of the genome according to RepeatMasker and 42% of the genome based on RepeatExplorer analysis.

In addition to characterizing the *G. anomalum* genome via clustering, we also co-clustered a diverse array of previously sequenced species (see methods) to evaluate the repeat content of *G. anomalum* in the broader context of the genus (Supplementary Table 1). This clustering included at least one member of each lettered cotton “genome group” (i.e., A-G and K; Wang *et al*. 2018), which were all sampled to represent 1% of their genome size (Hendrix and Stewart 2005). Principal components analysis (PCA; Figure 2) generally separates the species by geography on the first axis, with the American “D-genome” cottons toward the left part of the plot, the Australian species (groups C, G, and K) toward the right, and the African species (genome groups A, B, E, and F) intermediate between those two. Notably, the PCA groupings loosely follow the phylogenetic relationships among the genome groups (Cronn *et al*. 2002). Relative to other cotton species, *G. anomalum* (B-genome) has an intermediate amount of TEs (Figure 3; Table 3), as expected from its intermediate genome size. Like most of the other cottons, the *G. anomalum* genome consists of approximately half repetitive sequences (627 Mb), most of which (90%) are *gypsy* elements.

**Figure 2.**
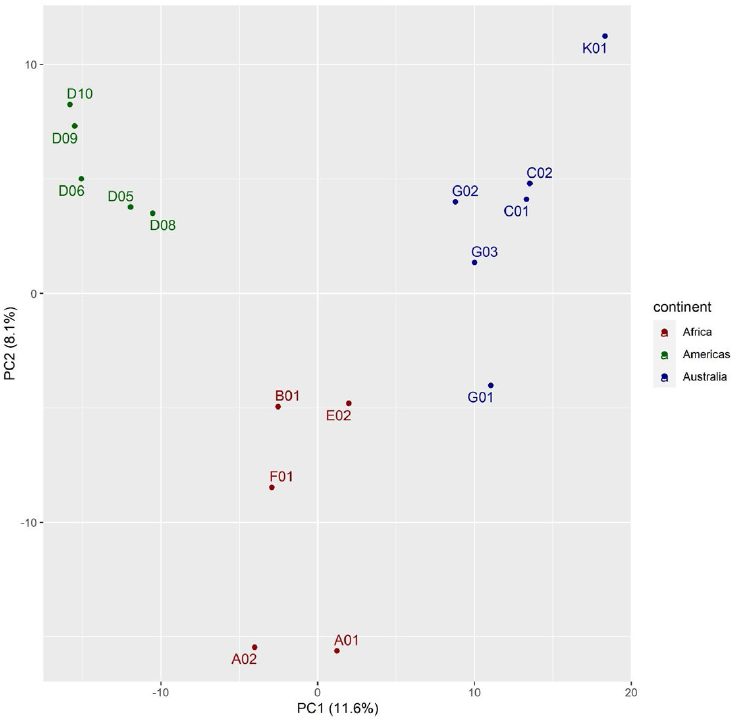
PCA analysis of repeats in cotton species. Species placement on the first two axes is primarily due to a small number of *gypsy* clusters. Species are colored by their broad geographic groups, i.e., the Americas (green), Africa/Arabian Peninsula (red), and Australia (blue) and listed by their official designations (Wang *et al*. 2018). The American cottons are *G. raimondii* (D5), *G. gossypioides* (D6), *G. trilobum* (D8), *G. laxum* (D9), and *G. turneri* (D10). The African/Arabian cottons are *G. herbaceum* (A1), *G. arboreum* (A2), *G. anomalum* (B1), *G. somalense* (E2), and *G. longicalyx* (F1). The Australian cottons are *G. sturtianum* (C1), *G. robinsonii* (C2), *G. bickii* (G1), *G. australe* (G2), *G. nelsonii* (G3), and *G. exiguum* (K1).

**Figure 3.**
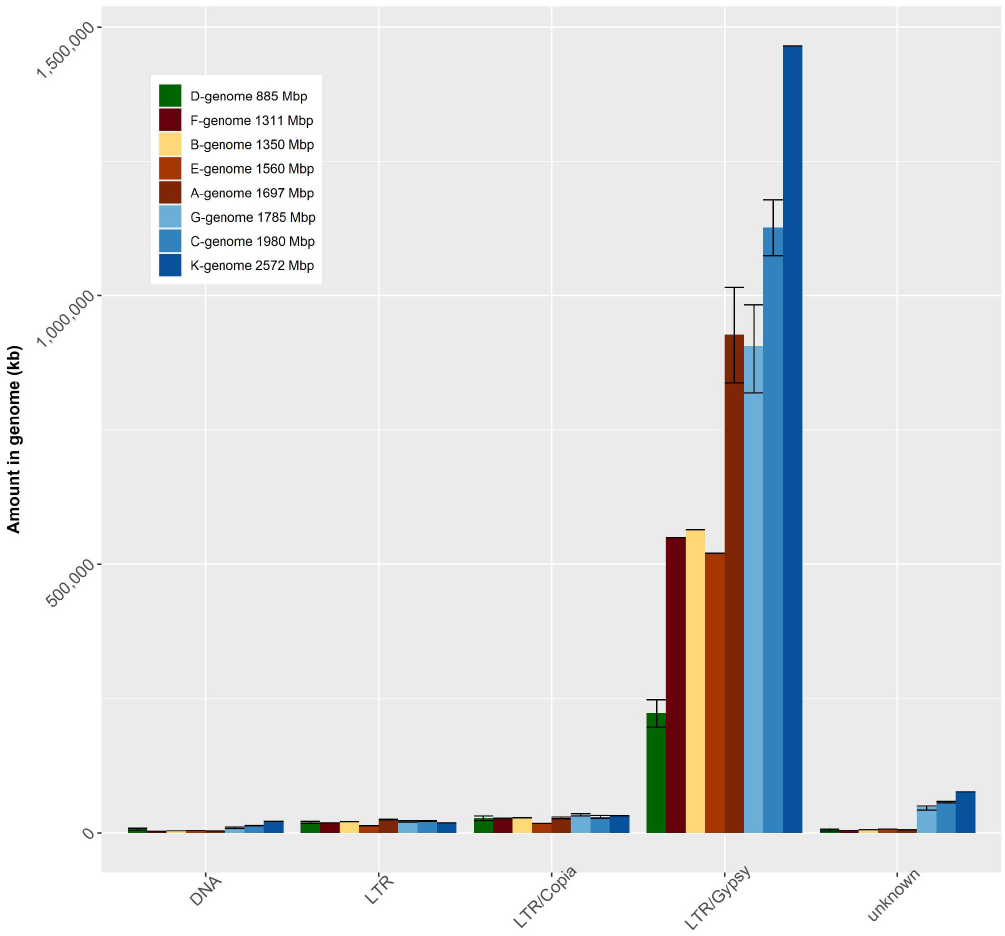
Average transposable element amounts in each genome group. Genome groups follow (Wang *et al*. 2018), and genome sizes for each genome group are from (Hendrix and Stewart 2005). Here, *G. anomalum* (yellow) is the sole representative of the B-genome clade.

**Table 3.** Repetitive amounts (average) in *Gossypium* genome groups compared to *G. anomalum* (B). Species representing each genome group are given in Supplementary Table 1.

| Genome Group | Geographic location | Genome Size (Mb) | Repeats (Mb) | Repeats (%) | Gypsy (Mb) | Gypsy (% repeats) | Gypsy (% genome) |
| --- | --- | --- | --- | --- | --- | --- | --- |
| **B** | **Africa** | **1350** | **627.1** | **47%** | **565** | **90%** | **42%** |
| A | Africa | 1697 | 992.9 | 59% | 927 | 93% | 55% |
| C | Australia | 1980 | 1253 | 63% | 1127 | 90% | 57% |
| D | Americas | 885 | 286.6 | 32% | 224 | 78% | 25% |
| E | Africa | 1560 | 567.5 | 36% | 520 | 92% | 33% |
| F | Africa | 1311 | 607.4 | 46% | 550 | 91% | 42% |
| G | Australia | 1785 | 1022 | 57% | 906 | 89% | 51% |
| K | Australia | 2572 | 1617.6 | 63% | 1465 | 91% | 57% |

Interestingly, while the *G. anomalum* genome is around 200 Mbp smaller than the African E-genome species (represented here by *G. somalense*), cluster analysis suggests that it has about 60 Mbp *more* repetitive sequences, most of which (40 Mb) are annotated as *gypsy* elements (Table 3). Regression analysis suggests that the amount of repetitive sequence observed in the E-genome clade is lower than expected, given the rest of the genome groups (Supplementary Figure 1). This may indicate general degradation and/or divergence in the repeats found in the E-genome clade, possibly indicating the presence of older elements, and/or that prior estimates of genome size are overestimates. The latter hypothesis would be consistent with the high contiguity and quality of our assembly that nevertheless recovered only 88% of the expected genome size.

Also notable is the observation that while the A-genome species (represented by both extant species, *G. herbaceum* and *G. arboreum*) are only ∼350 Mbp larger than *G. anomalum*, clustering suggests that they have approximately 1.5x more repetitive sequences, mostly *gypsy* (927 Mbp in A, versus 565 Mbp in B). This is, however, within what is expected as genome sizes in *Gossypium* increase (Supplementary Figure 1).

### *G. anomalum* as a vehicle to understand hybrid lethality in cotton

Hybrid lethality is a postzygotic reproductive barrier that results in embryo and/or seedling death in crosses involving incompatible plants, resulting in reduction and/or elimination of gene flow between populations or species (Bomblies and Weigel 2007; Maheshwari and Barbash 2011). While interspecific incompatibilities are common between species from different genome groups in *Gossypium*, interfertility is quite common between species from the same genome group (Hutchinson 1932; Silow 1941; Stephens 1946; Gerstel 1954; Menzel and Brown 1955; Phillips and Merritt 1972; Phillips and Reid 1975; Lee 1981c). For example, crosses are possible for most combinations of the 14 recognized D-genome diploids, insofar as these have been tested. An exception to this generality involves hybrid lethality in crosses that involve species from subsection *Integrifolia* (i.e., *G. davidsonii* and *G. klotzschianum*; D3d and D3k, respectively). These sister species are incompatible with nearly every other species in the genus, with the exception of *G. longicalyx* (F-genome) and *G. anomalum* (Phillips 1963). Notably, in some cases this lethality can be circumvented by increasing germination and growth temperatures (Phillips 1977), making lethality potentially useful in cultivar development (Lee 1981a).

While loci conferring hybrid lethality have been genetically identified through crosses and/or hexaploid bridging (Lee 1981a, 1981c; Endrizzi *et al*. 1985; Stelly 1990; Samora *et al*. 1994; Song *et al*. 2009), the underlying gene(s) controlling the D3 incompatibility are not yet known. In the late 1970s to early 1980s, Joshua A Lee generated a synthetic 2(A2D3) allotetraploid (Supplemental Figure 2) as described above, using the trick that *G. anomalum* was apparently “null” for the incompatibility factor and thus could be introgressed into A2 for purposes of creating the novel allopolyploid. Using a scheme of repeated backcrossing into *G. arboreum* and testing for fertility with *G. davidsonii*, crosses were continued for an unknown number of generations, but until hybrid progeny were uniformly healthy. Thus, the interspecific F1 hybrids were really tri-species constructs, in part, containing an introgressed locus (or loci) from *G. anomalum* that permits crosses with D3 to survive; ostensibly, this locus codes for a lethality factor in wildtype D3. Progeny from the last successful *G. arboreum* (BC) x *G. davidsonii* was subsequently doubled to create the synthetic 2(A2D3). This synthetic allotetraploid is thus primarily composed of *G. arboreum* and *G. davidsonii*, containing only a residual contribution from *G. anomalum*.

At present, the nature of the gene or genes controlling this hybrid lethality are unknown. Previous cytogenetic work on D3-lethality suggests that a single locus in *G. davidsonii* (i.e., Le^dav^) is responsible for lethality (Lee 1981b), and that this may interact with 1-2 loci in other cotton species (Lee 1981b; Stelly 1990). We downloaded resequencing reads from 2(A2D3) and a second synthetic allotetraploid (i.e., 2(A2D1)), which is a doubled *G. arboreum* x *G. thurberi* (Beasley 1940), and thus similar to 2(A2D3) but lacking the *G. anomalum* introgression. Competitive mapping of both synthetic allotetraploids to a reference containing the combined genomes of *G. arboreum, G. anomalum*, and *G. raimondii* (i.e., hereafter ABD-reference) reveals that approximately 1-2% of reads in each synthetic map strictly to *G. anomalum* chromosomes (Table 4), with a slightly higher percentage of reads from 2(A2D3) characterized as B-like (1.97 versus 1.66%). The number of reads considered A-or D-like is over an order of magnitude higher for both synthetics. Reads that could not be distinguished as A-, B-, or D-like (due to shared ancestry) were discarded from all samples, retaining approximately 70-75% of the reads. Because symplesiomorphy, autapomorphy, and technical error all have the potential to confound species identification of reads, we filtered locations in the ABD-reference *G. anomalum* chromosomes where we unexpectedly observed mapping of *G. arboreum, G. davidsonii*, and/or 2(A2D1)-derived reads, all of which should not have a *G. anomalum* origin. The remaining regions were considered markers for candidate locations where *G. anomalum* introgression remains in the 2(A2D3) synthetic allotetraploid.

**Table 4.** Reads (in million; M) uniquely mapped to *G. arboreum* (A2), *G. anomalum* (B1), and *G. raimondii* (D5).

| Species | total | mapped | B1 | B1 % | A2 | A2 % | D5 | D5 % |
| --- | --- | --- | --- | --- | --- | --- | --- | --- |
| *G. davidsonii* | 242.34 | 240.32 | 7.27 | 3.0 | 10.12 | 4.2 | 148.05 | 61.6 |
| *G. davidsonii* | 337.44 | 334.70 | 8.62 | 2.6 | 9.09 | 2.7 | 196.92 | 58.8 |
| 2(A2D1) | 310.72 | 295.84 | 4.90 | 1.7 | 159.14 | 53.8 | 59.25 | 20.0 |
| 2(A2D3) | 322.04 | 316.19 | 6.23 | 2.0 | 132.88 | 42.0 | 80.05 | 25.3 |
| *G. anomalum* | 305.98 | 297.64 | 250.27 | 84.1 | 5.23 | 1.8 | 0.72 | 0.2 |
| *G. anomalum* | 269.21 | 263.53 | 221.04 | 83.9 | 4.62 | 1.8 | 0.64 | 0.2 |
| *G. anomalum* | 132.07 | 128.74 | 107.68 | 83.7 | 2.35 | 1.8 | 0.32 | 0.3 |
| *G. anomalum* | 219.44 | 218.38 | 183.82 | 84.2 | 7.06 | 3.2 | 0.97 | 0.4 |
| *G. arboreum* | 478.36 | 476.17 | 5.46 | 1.2 | 346.42 | 72.8 | 1.72 | 0.4 |
| *G. arboreum* | 424.56 | 422.36 | 4.77 | 1.1 | 308.67 | 73.1 | 1.60 | 0.4 |
| *G. arboreum* | 415.34 | 414.20 | 4.49 | 1.1 | 302.71 | 73.1 | 1.29 | 0.3 |

After discarding short (<5 kb) regions as putative artifacts, we identified 28 regions on 9 chromosomes with putative introgression (Supplementary Table 5), for a total length of 195.7 kb. Most chromosomes exhibit small, discontiguous regions of putative introgression (<25 kb in length); however, a 287.5 kb window on chromosome B06 contains 13 of the 28 regions that collective comprise 69% (111.5 kb) of the total introgressed *G. anomalum* sequence. Genome annotation in this putative introgressed hotspot reveals only two gene models (i.e., B06G223600 and B06G223900) that overlap with the B-like regions, suggesting that one or both of these genes may be important for conferring fertility with *Integrifolia* (D3) species. Although these gene models are near-sequential in the genome, they are separated by over 172 kb of intervening sequence, as well as two additional genes contained within the 287.5 kb window that do not exhibit evidence of introgression. The first gene, B06G223600, is a putative F-box/kelch-repeat protein similar to At4g19870, whereas the second (B06G223900) is similar to PAP12, a phosphatase from *Arabidopsis thaliana*. Notably, in 2(A2D3), B06G223600 has 15 bp of extra sequence relative to the A-genome ortholog, representing an additional 5 amino acids in the protein. The second gene, B06G223900, however has no obvious sequence or structural differences, other than increased heterozygosity representing the presence of both A-and B-derived alleles. Further research, including expression-based analyses, will be required to fully understand the contribution of these and/or other genes to D3-lethality in cotton.

## Conclusion

The cotton genus has been the beneficiary of multiple high-quality genome sequences. While many have focused on the domesticated species, recent efforts have led to the generation of reference genomes for some of the wild representatives among the approximately 50 species in the genus (Cai *et al*. 2019; Grover *et al*. 2020, 2021; Chen *et al*. 2020; Wang *et al*. 2021). Here we report the first *de novo* sequence for a representative of the B-genome (Wang *et al*. 2018), whose members provide additional germplasm resources for both understanding and incorporating features like stress resistance and/or hybrid lethality into breeding programs. This resource will provide the foundation for future research into cotton diseases, such as blackarm (Knight 1954; Fryxell *et al*. 1984), as well as provide a potential source for fiber quality improvements (Mehetre 2010) and/or fertility control among different cotton lines.

## Acknowledgements

We thank and remember the late Joshua A Lee for his contributions to science, including improving our understanding of hybrid lethality and for creating the 2(A2D3) synthetic. We thank the National Science Foundation Plant Genome Research Program (Grant #1339412), the United States Dept. of Agriculture -Agriculture Research Service (Grant #58-6066-6-046), and Cotton Inc. for their financial support. We thank the Iowa State University ResearchIT unit, the BYU Fulton SuperComputer lab, the USDA-ARS, and the Mississippi State University High Performance Computing Collaboratory for computational resources and support.

---

**Supplementary Figure 1:**
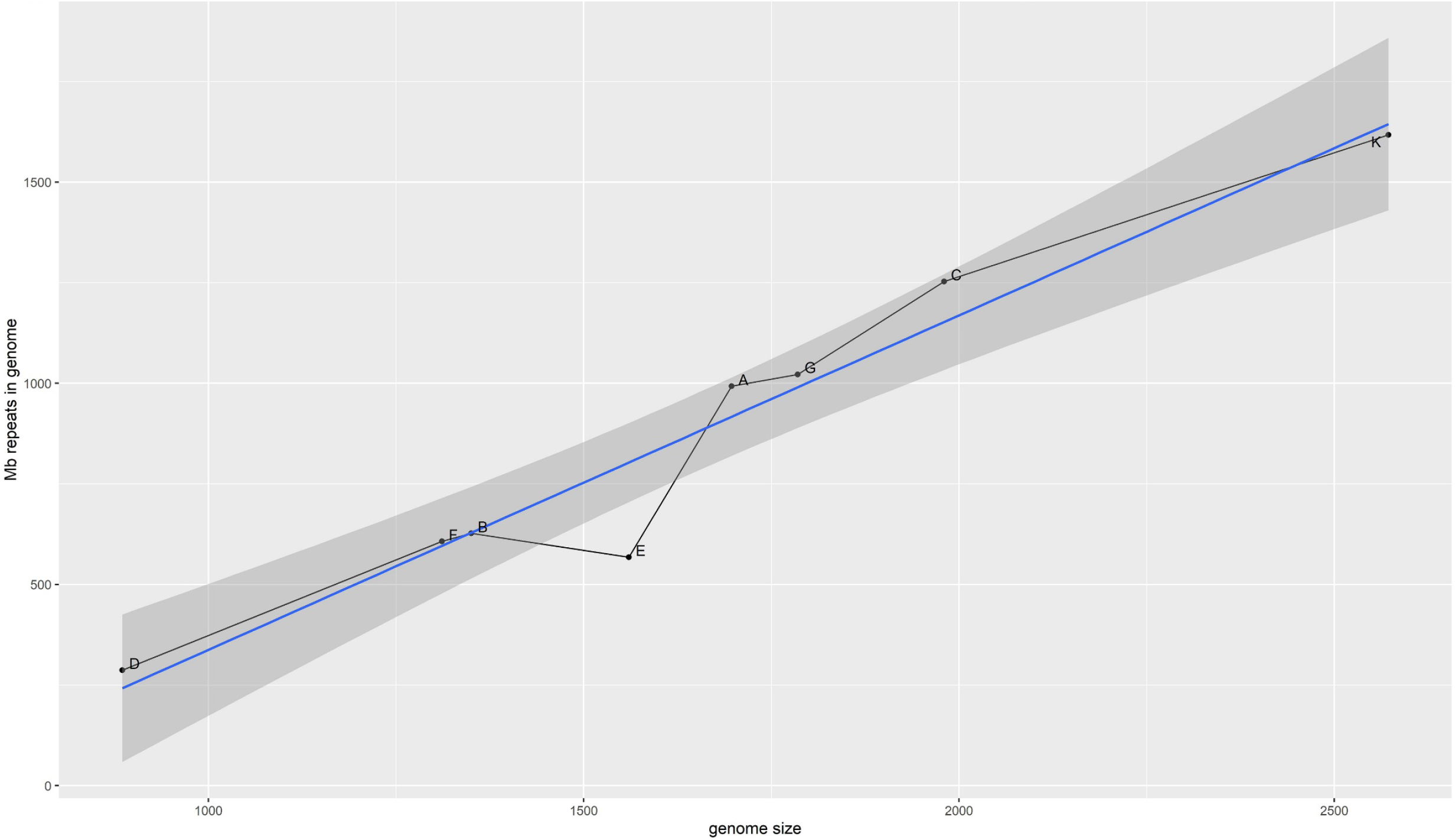
Genome size (Mbp) versus the amount of predicted repetitive sequence (in Mbp).

**Supplementary Figure 2:**
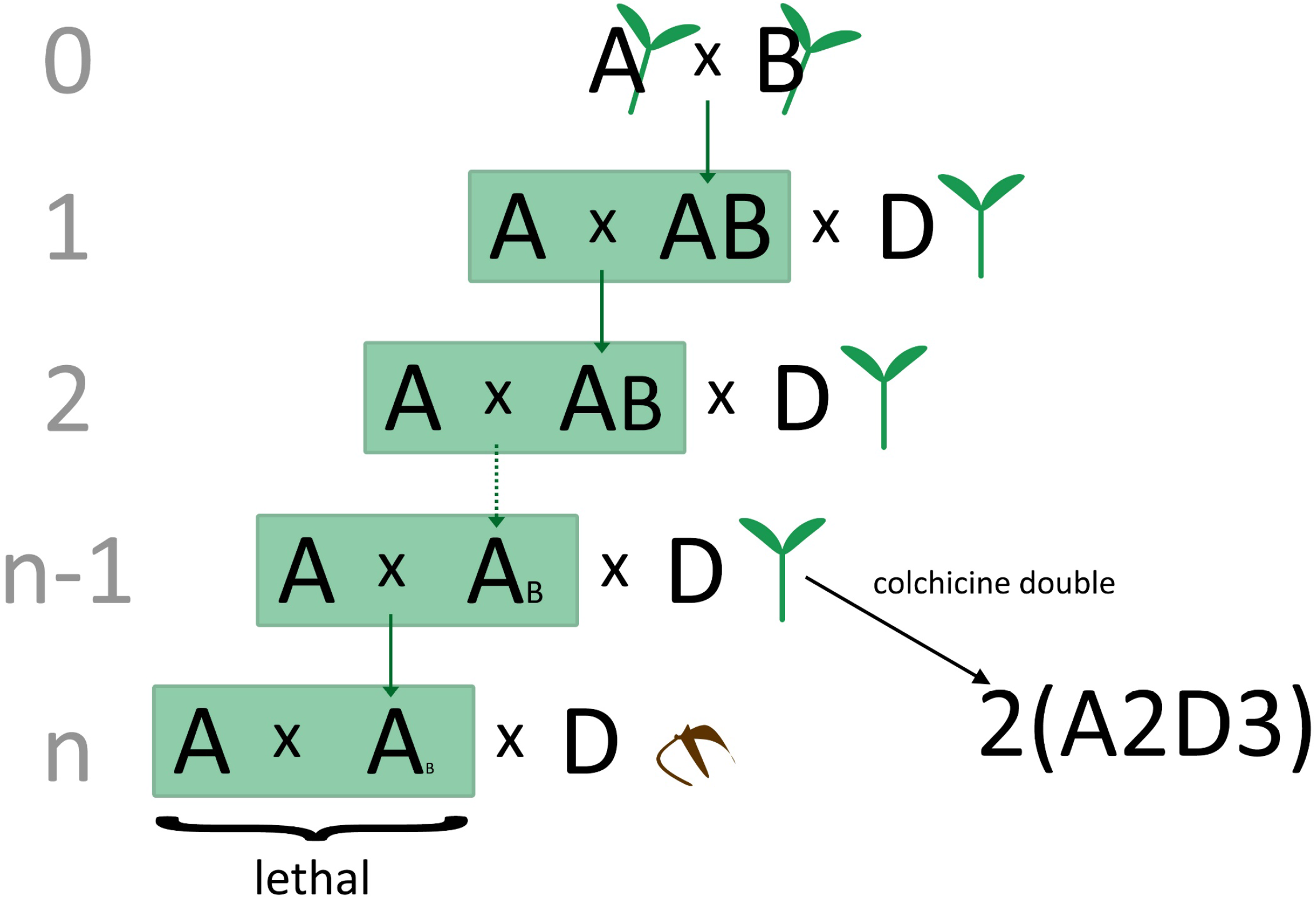
Diagram of the crossing scheme used to create the synthetic 2(A2D3). Crosses with *G. davidsonii* (D) test the viability of the introgressed *G. arboreum* line with *G. davidsonii*. Backcrossing with *G. arboreum* was continued until fertility was lost. The last fertile lineage was crossed with *G. davidsonii* and subsequently doubled to make 2(A2D3).

**Supplementary Table 1.** Genomic sequences from SRA used in *G. anomalum* analyses.

| **Species** | **Genome code** | **SRA number** | **Method** |
| --- | --- | --- | --- |
| 2(A2D3) | 2(A2D3) | SRR6334602 | genome mapping |
| 2(A2D3) | 2(A2D3) | SRR6334601 | genome mapping |
| *Gossypium anomalum* | B01 | SRR3560153 | genome mapping |
| *Gossypium anomalum* | B01 | SRR3560155 | genome mapping |
| *Gossypium anomalum* | B01 | SRR3560156 | genome mapping |
| *Gossypium anomalum* | B01 | SRR12745560 | genome mapping |
| *Gossypium arboreum* | A02 | SRR8979965 | genome mapping |
| *Gossypium arboreum* | A02 | SRR8979944 | genome mapping |
| *Gossypium arboreum* | A02 | SRR8979925 | genome mapping |
| *Gossypium davidsonii* | D03 | SRR6334584 | genome mapping |
| *Gossypium davidsonii* | D03 | SRR8136261 | genome mapping |

| **Species** | **Genome code** | **SRA number** | **Method** | **Genome size** |
| --- | --- | --- | --- | --- |
| *G. herbaceum* | A01 | SRR8979969 | clustering | 1697 |
| *G. arboreum* | A02 | SRR8979922 | clustering | 1710 |
| *G. anomalum* | B01 | SRR3560153 | clustering | 1359 |
| *G. sturtianum* | C01 | SRR8979990 | clustering | 2015 |
| *G. robinsonii* | C02 | SRR8979901 | clustering | 1951 |
| *G. raimondii* | D05 | SRR847980 | clustering | 880 |
| *G. gossypoides* | D06 | SRR3560149 | clustering | 841 |
| *G. trilobum* | D08 | SRR8136271 | clustering | 851 |
| *G. laxum* | D09 | SRR8136274 | clustering | 934 |
| *G. turneri* | D10 | SRR8136255 | clustering | 910 |
| *G. somalense* | E02 | SRR3560162 | clustering | 1496 |
| *G. longicalyx* | F01 | SRR617704 | clustering | 1311 |
| *G. bickii* | G01 | SRR3560189 | clustering | 1756 |
| *G. australe* | G02 | SRR8979992 | clustering | 1834 |
| *G. nelsonii* | G03 | SRR8979903 | clustering | 1756 |
| *G. exiguum* | K01 | SRR3560141 | clustering | 2460 |

**Supplementary Table 2.**
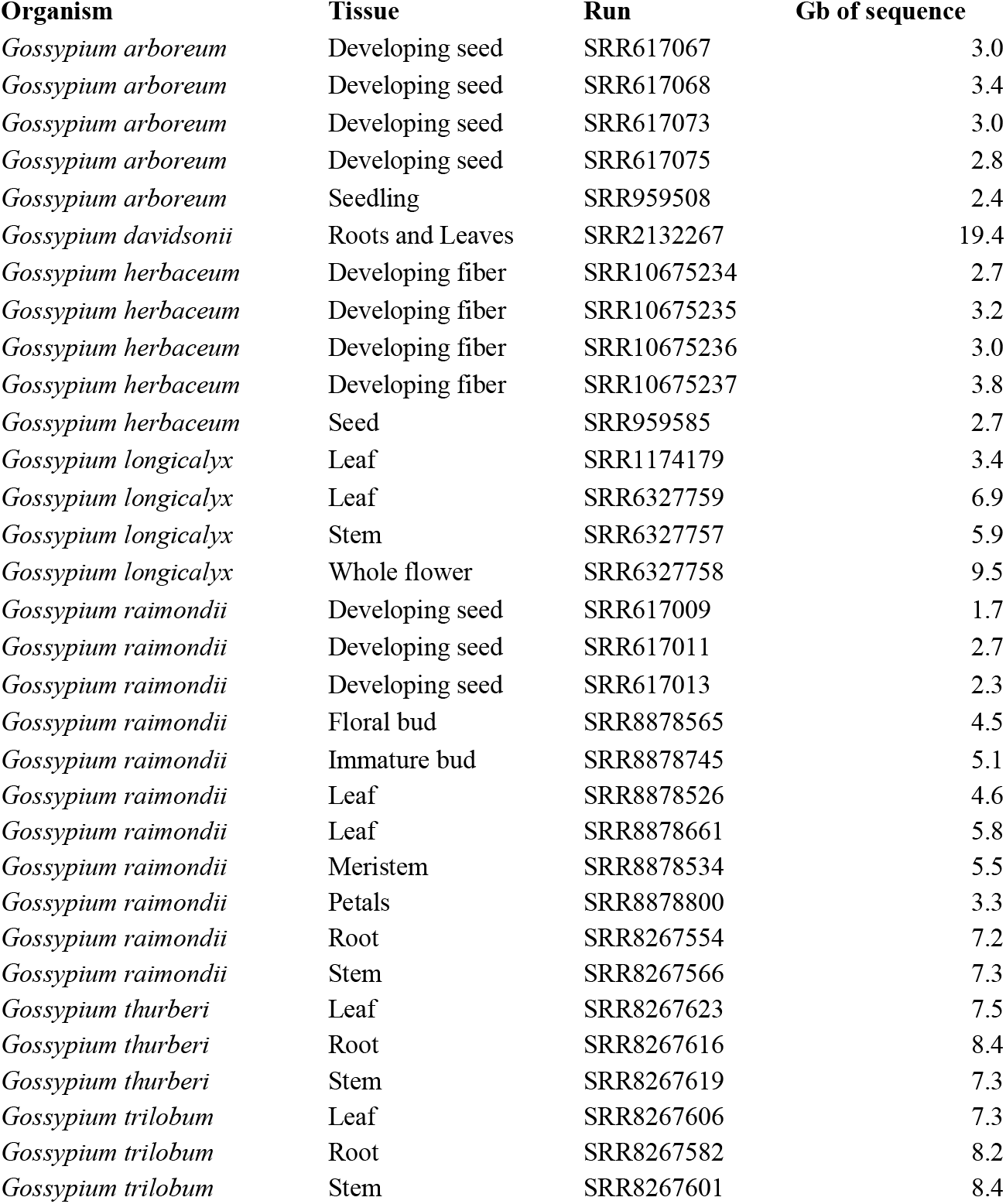
RNA-seq downloaded from the SRA and used to annotate the *G. anomalum* genome.

| **Organism** | **Tissue** | **Run** | **Gb of sequence** |
| --- | --- | --- | --- |
| *Gossypium arboreum* | Developing seed | SRR617067 | 3.0 |
| *Gossypium arboreum* | Developing seed | SRR617068 | 3.4 |
| *Gossypium arboreum* | Developing seed | SRR617073 | 3.0 |
| *Gossypium arboreum* | Developing seed | SRR617075 | 2.8 |
| *Gossypium arboreum* | Seedling | SRR959508 | 2.4 |
| *Gossypium davidsonii* | Roots and Leaves | SRR2132267 | 19.4 |
| *Gossypium herbaceum* | Developing fiber | SRR10675234 | 2.7 |
| *Gossypium herbaceum* | Developing fiber | SRR10675235 | 3.2 |
| *Gossypium herbaceum* | Developing fiber | SRR10675236 | 3.0 |
| *Gossypium herbaceum* | Developing fiber | SRR10675237 | 3.8 |
| *Gossypium herbaceum* | Seed | SRR959585 | 2.7 |
| *Gossypium longicalyx* | Leaf | SRR1174179 | 3.4 |
| *Gossypium longicalyx* | Leaf | SRR6327759 | 6.9 |
| *Gossypium longicalyx* | Stem | SRR6327757 | 5.9 |
| *Gossypium longicalyx* | Whole flower | SRR6327758 | 9.5 |
| *Gossypium raimondii* | Developing seed | SRR617009 | 1.7 |
| *Gossypium raimondii* | Developing seed | SRR617011 | 2.7 |
| *Gossypium raimondii* | Developing seed | SRR617013 | 2.3 |
| *Gossypium raimondii* | Floral bud | SRR8878565 | 4.5 |
| *Gossypium raimondii* | Immature bud | SRR8878745 | 5.1 |
| *Gossypium raimondii* | Leaf | SRR8878526 | 4.6 |
| *Gossypium raimondii* | Leaf | SRR8878661 | 5.8 |
| *Gossypium raimondii* | Meristem | SRR8878534 | 5.5 |
| *Gossypium raimondii* | Petals | SRR8878800 | 3.3 |
| *Gossypium raimondii* | Root | SRR8267554 | 7.2 |
| *Gossypium raimondii* | Stem | SRR8267566 | 7.3 |
| *Gossypium thurberi* | Leaf | SRR8267623 | 7.5 |
| *Gossypium thurberi* | Root | SRR8267616 | 8.4 |
| *Gossypium thurberi* | Stem | SRR8267619 | 7.3 |
| *Gossypium trilobum* | Leaf | SRR8267606 | 7.3 |
| *Gossypium trilobum* | Root | SRR8267582 | 8.2 |
| *Gossypium trilobum* | Stem | SRR8267601 | 8.4 |

**Supplementary Table 3.** *G. anomalum*-specific orthogroups. Orthogroups are from Grover et al (2021).

|  |  |  |  |  |
| --- | --- | --- | --- | --- |
| OG0028671 | Goano.001G191300 | Goano.003G147100 | Goano.003G220400 | Goano.010G245100 |
| OG0030787 | Goano.003G136200 | Goano.009G026000 | Goano.011G329200 |  |
| OG0034106 | Goano.010G047200 | Goano.010G047400 |  |  |

**Supplementary Table 4.**
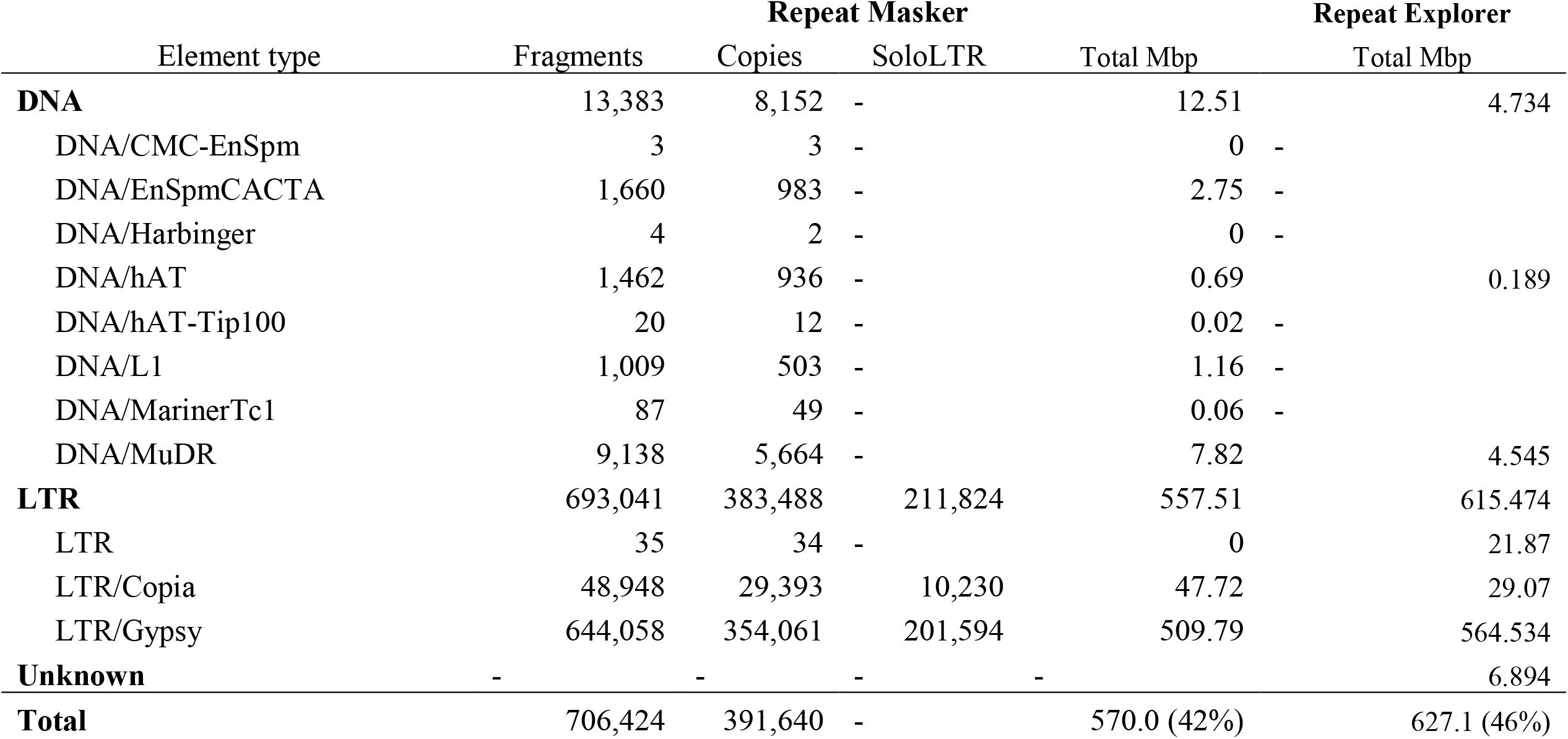
Comparison between RepeatMasker and RepeatExplorer outputs for the *G. anomalum* genome.

**Supplementary Table 5.** Regions of putative *G. anomalum* introgression in 2(A2D3) and gene models within those regions.

| Chromosome | Start | Stop | Length | Chromosome total (bp) | Gene | Gene Start | Gene Stop | Function |
| --- | --- | --- | --- | --- | --- | --- | --- | --- |
| B_02 | 1,751,517 | 1,759,641 | 8,124 | 19,076 | B1.B_02G017200 | 1,759,353 | 1,759,641 | NGA4: B3 domain-containing transcription factor NGA4 |
| B_02 | 20,496,842 | 20,502,002 | 5,160 |  |  |  |  |  |
| B_02 | 70,412,653 | 70,418,445 | 5,792 |  |  |  |  |  |
| B_03 | 106,013,266 | 106,019,946 | 6,680 | 6,680 | B1.B_03G260300 | 106,013,267 | 106,019,946 | Os03g0733400: Zinc finger BED domain-containing protein RICESLEEPER 2 |
| B_04 | 50,725,528 | 50,730,720 | 5,192 | 5,192 |  |  |  |  |
| B_05 | 2,823,728 | 2,829,382 | 5,654 | 10,948 |  |  |  |  |
| B_05 | 10,464,363 | 10,469,657 | 5,294 |  | B1.B_05G053800 | 10,464,364 | 10,465,910 | At3g47570: Probable LRR receptor-like serine/threonine-protein kinase At3g47570 |
| B_06 | 92,554,341 | 92,562,783 | 8,442 | 111,544 |  |  |  |  |
| B_06 | 92,562,858 | 92,570,652 | 7,794 |  | B1.B_06G223600 | 92,567,389 | 92,568,340 | At4g19870: F-box/kelch-repeat protein At4g19870 |
| B_06 | 92,641,196 | 92,646,951 | 5,755 |  |  |  |  |  |
| B_06 | 92,646,956 | 92,657,533 | 10,577 |  |  |  |  |  |
| B_06 | 92,670,418 | 92,675,475 | 5,057 |  |  |  |  |  |
| B_06 | 92,675,847 | 92,681,845 | 5,998 |  |  |  |  |  |
| B_06 | 92,681,890 | 92,694,410 | 12,520 |  |  |  |  |  |
| B_06 | 92,736,613 | 92,744,234 | 7,621 |  | B1.B_06G223900 | 92,741,077 | 92,741,403 | PAP12: Fe(3+)-Zn(2+) purple acid phosphatase 12; |
| B_06 | 92,748,186 | 92,760,871 | 12,685 |  |  |  |  |  |
| B_06 | 92,774,479 | 92,782,790 | 8,311 |  |  |  |  |  |
| B_06 | 92,785,370 | 92,790,493 | 5,123 |  |  |  |  |  |
| B_06 | 92,819,750 | 92,830,095 | 10,345 |  |  |  |  |  |
| B_06 | 92,830,555 | 92,841,871 | 11,316 |  |  |  |  |  |
| B_07 | 76,245,298 | 76,251,283 | 5,985 |  |  |  |  |  |
| B_08 | 15,137,345 | 15,143,614 | 6,269 | 12,335 |  |  |  |  |
| B_08 | 94,864,034 | 94,870,100 | 6,066 |  | B1.B_08G233100 | 94,864,035 | 94,870,100 | HMA5: Probable copper-transporting ATPase HMA5 |
| B_11 | 51,368,114 | 51,373,226 | 5,112 | 24,037 |  |  |  |  |
| B_11 | 86,773,815 | 86,779,971 | 6,156 |  |  |  |  |  |
| B_11 | 95,280,456 | 95,287,671 | 7,215 |  | B1.B_11G340700 | 95,280,537 | 95,281,955 | Retrovirus-related Pol polyprotein from transposon TNT 1-94 |
| B_11 | 99,060,757 | 99,066,311 | 5,554 |  | B1.B_11G362400 | 99,060,758 | 99,066,311 | At3g47200: UPF0481 protein At3g47200; |
| B_12 | 17,145,074 | 17,150,915 | 5,841 | 5,841 |  |  |  |  |

